# Network-enabled drug repurposing identifies lithium to promote cardiomyocyte proliferation

**DOI:** 10.64898/2026.10.15.759417

**Authors:** Kaitlyn L. Wintruba, Sunil Verma, Irfan S. Kathiriya, Aleksandra Kostina, Aitor Aguirre, Muge Kuyumcu-Martinez, Jeffrey J. Saucerman

## Abstract

The adult mammalian heart has minimal regenerative capacity, limited in part by endogenous cardiomyocyte (CM) renewal. Although CM proliferation is a promising target in both congenital and adult heart disease, no current therapies specifically target this process. Progress has been hindered by the complexity of CM proliferation signaling networks, which may require systems-level approaches. The overall goal of this study was to leverage systems modeling to identify FDA approved drugs that have the potential to be repurposed towards CM proliferation. To achieve this goal, we developed a computational drug repurposing framework that integrates drug-target interaction data with a previously developed logic-based network model of CM proliferation signaling. Using this framework, we identified 81 drugs that directly target nodes within the CM proliferation network, corresponding to 32 unique drug-target interactions. Virtual screening predicted 5 unique drug-target interactions that increase CM proliferation and 14 that decrease CM proliferation outputs. Lithium, a GSK3β inhibitor widely prescribed for bipolar disorder, was predicted and experimentally validated to promote CM proliferation through a Cyclin D/CDK4 pathway. *In vivo*, treatment of pregnant mice with lithium increased DNA synthesis in the CMs of their embryos. A retrospective analysis of the FDA Adverse Events Reporting System confirmed that relative to other mood stabilizers, *in utero* exposure to lithium was associated with an increased incidence of congenital heart defects, in males but not females. Together, this work provides a computational-experimental framework for repurposing FDA-approved drugs to modulate CM proliferation, which highlights therapeutic potential of lithium for adult CM regeneration but congenital risks to developing embryos.

## Introduction

Heart disease remains a leading cause of morbidity and mortality worldwide^1^. A central driver of this burden is the irreversible loss of functional cardiomyocytes (CMs) following cardiac injury, a consequence of the limited proliferative capacity of adult human CMs^2^. During embryonic development, CM proliferation is tightly regulated and persists for only a brief postnatal window before the myocardium transitions to a largely post-mitotic state^3^. In the context of congenital heart disease, this critical developmental transition is often disrupted, impairing the CM proliferation necessary for normal cardiac growth and repair, though the mechanisms underlying this disruption remain incompletely understood^4^. Therefore, stimulating endogenous CM proliferation represents a compelling therapeutic strategy with relevance to both adult and congenital heart disease, where restoration of CM number could meaningfully improve cardiac function.

Studies using genetic approaches have demonstrated that CM proliferation can be reactivated to improve outcomes in a variety of adult cardiac disease models. Overexpression of transcription factors such as Foxf1/2^5^, and Myc^6^, as well as activation of the Hippo-YAP signaling pathway^7,8^ have each shown promising regenerative effects in preclinical models. However, genetic manipulation strategies face translational barriers such as high cost, need for specific delivery, and safety concerns, underscoring the need for pharmacological alternatives that can be safely used in patients.

Drug repurposing offers an attractive path forward. Drugs already approved by the FDA for other indications carry established safety profiles and known pharmacokinetics, substantially reducing the translational hurdles inherent to *de novo* drug development^9^. This shift toward human-relevant, non-animal approaches is increasingly supported by the FDA and NIH, which have prioritized new approach methodologies (NAMs), such as human iPSC-derived cardiomyocytes, cardiac organoids, and *in silico* cardiac models, to better predict human cardiac responses to therapeutics^10^.

While computational approaches have great promise for accelerating the drug development pipeline^11,12^, direct application to drugs that induce CM proliferation remain limited. A recent study used molecular docking simulations against specific known targets Meis1 and Hoxb13, and identified the aminoglycoside antibiotics paromomycin and neomycin as inducers of cardiac regeneration in cells and animal models^13^. Computational frameworks that integrate the broader CM cell cycle signaling network with simulations of drug responses may enable expansion of this paradigm to a wider range of targets and candidate drugs for experimental validation^14^.

In this study, we develop a virtual drug screening pipeline to identify FDA-approved drugs predicted to promote CM proliferation. Our approach maps FDA-approved drugs to protein targets that are mechanistically predicted to be influential in modulating phenotypes associated with CM proliferation. By applying this pipeline, we identify lithium as a top candidate for drug repurposing, and we validate its pro-proliferative effects using *in vitro* neonatal rat CMs, an *in vivo* mouse embryonic heart model, and clinical data.

## Methods

### Logic-based model of the cardiomyocyte proliferation signaling network

As a foundation for mechanistic network simulations, we used our recent logic-based network model of the CM proliferation signaling network^15^. This network model was manually constructed from published literature and integrated 72 nodes and 88 reactions to define five phenotypic outputs (DNA replication, mitosis, polyploid, cytokinesis, and binucleation) of CM cell cycle. Signaling dynamics were predicted by mapping this network model into logic-based differential equations, implemented in Netflux^16,17^. Previously, this network model accurately predicted outcomes for 74 out of 81 experiments reported in the literature independent from that used to build the model, achieving a predictive accuracy of approximately 91%. Further, the model predicted Myc as a mediator of YAP-induced CM proliferation, which was validated with *in vitro* and *in vivo* data^15^.

### Virtual drug screen

Drugs targeting proteins within the network model were identified using DrugBank^18^ (version 5.1.12), a database containing drug target data on approximately 4,500 FDA approved and 6,000 clinically investigational drugs. For each identified drug, the name, database ID, known molecular target(s), and corresponding drug action data were extracted using a Python script. This process yielded 482 drugs targeting nodes of the network model, of which 81 had annotated drug action (agonist or antagonist) information. Some of the 81 drugs shared the same network target and action, resulting in 32 unique drug target-action combinations. To simulate drug activity, each unique target-action combination was integrated into the network model by modifying the activation function of the corresponding target node(s)^19^. Simulations were conducted by first allowing the network model to reach a steady state under baseline conditions, followed by application of a saturating drug dose (reaction weight = 1). For each simulation, the fold change in node activity was calculated as the ratio of the activity level under drug treatment to the baseline level.

### Mechanistic subnetwork analysis

Mechanistic subnetworks are defined as the intersection between a) nodes directly affected by a given drug and b) nodes whose knockdown modulates the effect of a drug on a downstream phenotype^20^. To determine which nodes mediated the drug’s effects on the proliferation outputs, a sensitivity analysis was performed by systematically knocking down individual nodes by setting their respective Ymax parameter to 0 and running the simulation to steady state in the presence of the drug. Changes in network model outputs were used to identify nodes influential to the drug-induced response. These influential nodes were then compared to the set of nodes directly targeted or responsive to the drug. The overlapping nodes were used to create a mechanistic subnetwork, which was visualized in Cytoscape^21^ to identify the key components underlying the drug’s network mechanism of action.

### In vitro validation

All animal experiments were conducted in accordance with the NIH Guidelines for the Care and Use of Laboratory Animals and were approved by the University of Virginia Institutional Animal Care and Use Committee. CMs were isolated from 1-to 2-day-old Sprague-Dawley rats using the Neomyt isolation kit (Cellutron #NC9073658). Each isolation pooled CMs from all 9-12 male and female pups within a litter. Cells were plated at a density of 30,000 cells per well in 96-well Corning CellBIND plates pre-coated with SureCoat (Cellutron #NC9073658) and maintained in plating medium (low-glucose DMEM, 17% M199, 10% horse serum, 5% fetal bovine serum, 1% L-Glutamine, 10 U/mL penicillin, and 50 mg/mL streptomycin). After 48 hours of culture with daily media changes, cells were serum-starved with WE+B medium (William’s E medium without phenol red (Gibco #A1217601) supplemented with Primary Hepatocyte Cell Maintenance Supplement B (Gibco #CM4000) for 4 hours prior to treatment. CMs were then treated with individual drugs or drug combinations in serum-free media for 48 hours, with DMSO (0.1%) serving as the vehicle control.

Following treatment, cells were fixed in 4% paraformaldehyde and fluorescently labeled with anti-α-actinin (Sigma #A7811) or anti-cardiac troponin T (Abcam #ab45932), anti-Ki67

(Invitrogen #14-5698-82), anti-Cyclin D1 (abcam #ab16663), and DAPI (Sigma #D9542). Stained cells were imaged using the Operetta CLS high-content imaging system with a 10 × 0.3 NA objective. Images were analyzed using a CellProfiler^22^ pipeline for nuclear segmentation and quantification of Ki67 and Cyclin D1 expression in CMs as described in previous studies^15,23^. Briefly, nuclei were segmented from DAPI-stained images, and α-actinin or cardiac troponin T staining was used to identify nuclei belonging to CMs. Thresholds for mean nuclear Ki67 or Cyclin D1 intensity were set using triangle method^24^. Multinucleated CMs were identified by measuring the distance between adjacent nuclei, with a threshold of less than 2 pixels. DNA content was quantified from integrated DAPI intensities and combined with Ki67 positivity to classify nuclei into cell cycle phase and ploidy states as described in previous studies^25,26^.

### In vivo validation

Timed-pregnant C57BL/6 mice (Jackson Laboratory) were used for all experiments. Embryonic day (E) 0.5 was designated as noon on the day a vaginal plug was observed. Mice received a single intraperitoneal injection of either lithium carbonate (250 mg/kg) or saline (vehicle control) at E7.5. To label proliferating cells, 5-ethynyl-2′-deoxyuridine (EdU) (Invitrogen #E10187) was administered intraperitoneally 4 hours prior to embryo collection at a dose of 50 mg/kg. At E13.5, embryos were harvested following euthanasia of the dams by CO2 inhalation and cervical dislocation. Uteri were dissected and transferred to ice-cold phosphate-buffered saline (PBS), and embryonic hearts were isolated for downstream processing.

For each experimental group, 15 or 16 embryonic hearts were collected and fixed for 24 hours in 4% paraformaldehyde at 4°C. Hearts were rinsed in PBS, dehydrated through a graded ethanol series, cleared in xylene, and embedded in paraffin with proper orientation. Serial 5 μm-thick sections were cut using a microtome and mounted on positively charged glass slides (Fisher Scientific #22-037-246).

Following sectioning, embryonic heart sections were deparaffinized, subjected to antigen retrieval with 1X antigen unmasking solution (Vector Laboratories #H-3300-250), and processed for EdU detection using copper-catalyzed click chemistry solution (2 M Tris-HCl pH 8.5, 100 mM CuSO4, 10 µM 647 Azide (Fisher Scientific #10043964), and 250 mM ascorbic acid). Sections were then blocked and fluorescently labeled with anti-cardiac troponin T (cTnT, Abcam #ab45932). Slides were mounted using mounting medium containing DAPI (Sigma #F6057) and coverslipped before being imaged using the Leica THUNDER Imager with 20X objective. CM EdU incorporation in tissue sections stained for cTnT, EdU, and DAPI was quantified using Cellpose^27^(version 4.1.1, pretrained cpsam model) and ilastik^28^ (version 1.4.2). Nuclei were segmented from the DAPI channel of each 20X image tile in Cellpose. Segmented objects outside the tissue border were removed by manually outlining each section in napari, using a custom Python script. Nuclei masks and the corresponding stained images were then processed with the ilastik object classification workflow, in which manually annotated nuclei were used to train two separate random forest classifiers to label each nucleus as cTnT+ or cTnT- and as EdU+ or EdU-. Nuclei detected twice in the overlap between adjacent tiles were counted once. The percentage of EdU+ CM nuclei was calculated for each heart as the number of EdU+cTnT+ nuclei divided by the total number of cTnT+ nuclei.

### Retrospective analysis of pharmacovigilance data

Patient data were analyzed from the U.S. Food and Drug Administration’s Adverse Event Monitoring System (AEMS, formerly FAERS) database using the AERSmine tool^29^ and included reports from the first quarter of 2004 through the second quarter of 2024. AEMS reports were queried through AERSMine (version 2025-10-15, accessed June 11, 2026), and preferred terms were grouped by HLT, HLGT and SOC according to MedDRA (version 24.0). This retrospective study was exempt from informed consent because it involved secondary analysis of existing data, did not include interaction with human subjects, and used de-identified data in accordance with the HIPAA Privacy Rule (Section 164.514).

Children between the ages of 0 and 14 were analyzed to account for drug exposure occurring *in utero*. Congenital heart disease (CHD) was defined as 55 preferred terms: the 54 terms within the MedDRA high level group term “Congenital cardiac disorders” plus the term “Heart disease congenital”. Adult analyses used the MedDRA “Cardiomyopathies” and “Myocardial disorders NEC” high level terms (106 terms, covering cardiomyopathies, hypertrophy, systolic and diastolic dysfunction, fibrosis, and remodeling). across young adult, adult, and elderly age brackets.

For each drug, disproportionality was quantified as the reporting odds ratio (ROR) from a 2×2 contingency table comparing CHD or intrinsic myocardial disorder reports for that drug against all other reports in the database, with 95% Wald confidence intervals and Fisher’s exact test p-values corrected by the Benjamini-Hochberg method. A significant signal was defined as a 95% confidence interval lower bound > 1 with FDR q < 0.05 and at least 3 drug-event reports. Lithium was compared with the mood stabilizers valproic acid, carbamazepine, and lamotrigine.

### Statistics

Primary neonatal rat CMs were isolated from a minimum of two different isolations. For each isolation, the treatment groups had 3-5 replicate wells. Statistical significance of all experimental groups was determined by a two-way ANOVA (with isolation and treatment as groups) followed by Dunnett’s post-hoc test or Tukey’s multiple comparisons test. Embryonic mouse hearts were isolated from a minimum of 15 embryos per group and statistical significance determined by unpaired two-tailed Student’s t-test. For all analyses, a p-value <0.05 was considered statistically significant.

### Data availability

All data and code used in this manuscript are available on GitHub (https://github.com/saucermanlab/Wintruba-DrugRepurposing), made public with publication.

## Results

### Virtual drug screen predicts FDA-approved drugs that regulate cardiomyocyte proliferation

To identify putative therapeutics that modulate cardiomyocyte (CM) proliferation, we conducted a virtual drug screen by integrating systematic mapping of drugs and their targets to a recently developed mechanistic computational model of CM proliferation signaling^15^ (**Figure 1A**). This logic-based model of the CM proliferation signaling network was previously developed based on 23 papers, validated against 81 literature experiments with 91% accuracy, and applied to discover Myc as a driver of YAP-induced proliferation^15^. Here, we used the DrugBank database^18^ to identify 81 FDA-approved or investigational drugs that directly target nodes in the CM proliferation network model, which act either as agonists or antagonists (**Figure 1B**; full drug-target details in **Table S1**). For each of the 32 unique drug target-action combinations, we simulated their effects on the CM proliferation network activity (**Figure S1A**), revealing that the candidate drugs engage distinct intracellular pathways upstream of proliferative outputs.

**Figure 1.**
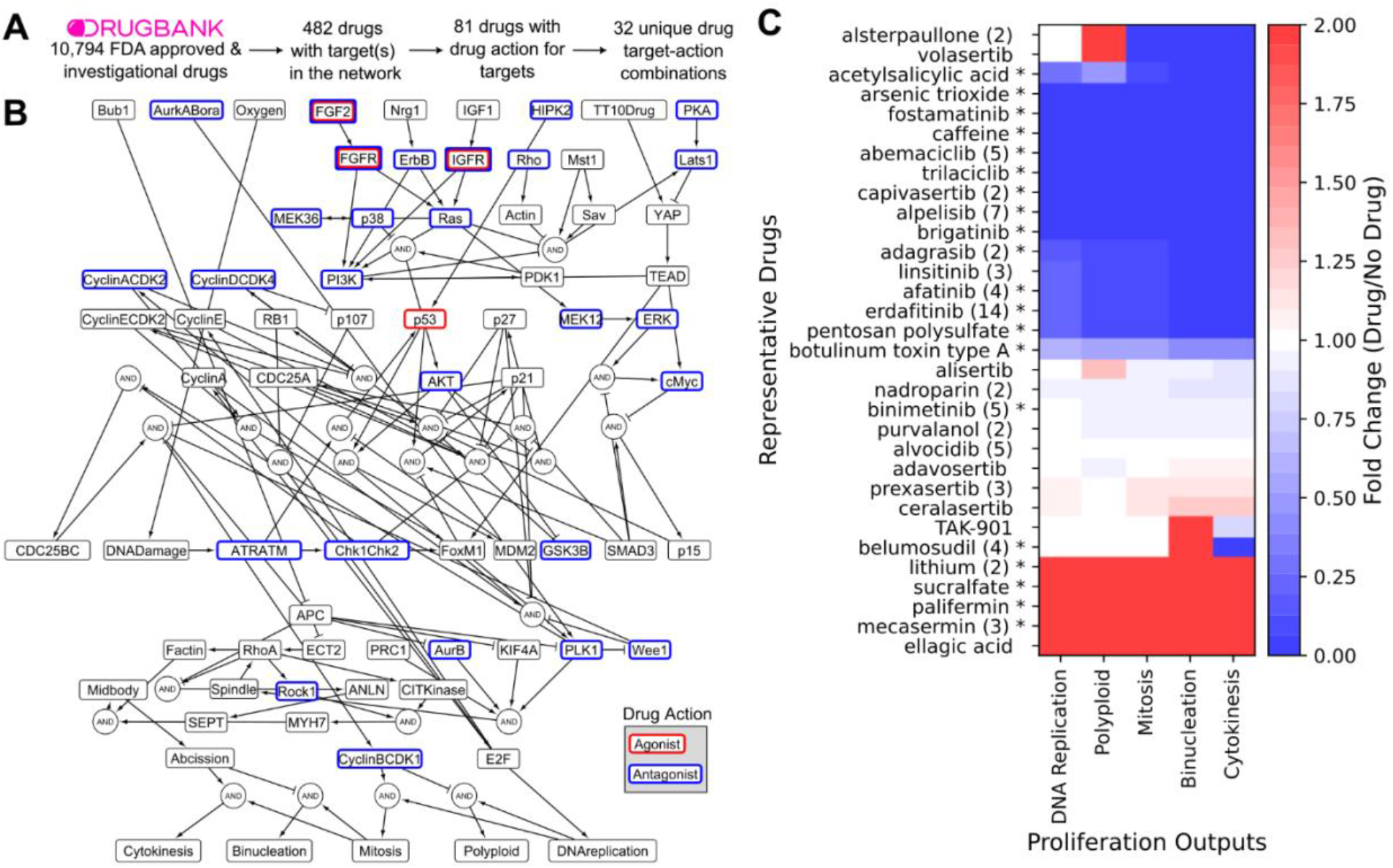
Virtual drug screen identifies drugs predicted to promote or suppress cardiomyocyte proliferation. **A)** Computational pipeline for identifying FDA approved and investigational drugs that target nodes in the network model. **B)** Network model of cardiomyocyte proliferation signaling, annotated based with drug targets according to the specified drug action. **C)** Network model simulations of representative drugs affecting proliferation outputs. (#) represents number of drugs exhibiting that unique drug target-action combination, and * indicates that the drug is FDA approved.

The virtual drug repurposing screen predicted the influence of each unique drug target-action combination on five proliferation-related outputs: DNA replication, mitosis, binucleation, cytokinesis, and polyploid. Among the 32 unique drug target-action combinations, 5 drugs (lithium, sucralfate, palifermin, mecasermin, and ellagic acid) were predicted to increase proliferation phenotypic outputs, while 15 were predicted to decrease them (**Figure 1C**). The remaining drugs either showed discordant effects across outputs or produced no significant changes (**Figure 1C**). Drugs with either pro- or anti-proliferative effects could conceivably modulate cardiac remodeling in a physiologic or pathologic manner^30^.

To prioritize candidates with translational potential for cardiac regeneration, we focused on three predicted pro-proliferative drugs with distinct molecular targets: lithium, an FDA-approved mood stabilizer that acts as a GSK3B inhibitor; sucralfate, an FDA-approved gastric ulcer treatment that acts as an FGF2 agonist; and ellagic acid, a naturally occurring polyphenol that acts as a PKA inhibitor. Dose-response simulations further distinguished their predicted effects across proliferative outputs (**Figure S1B**). Thus, these candidates were selected for experimental validation of their ability to promote CM proliferation *in vitro*.

**Supplementary Table 1.**
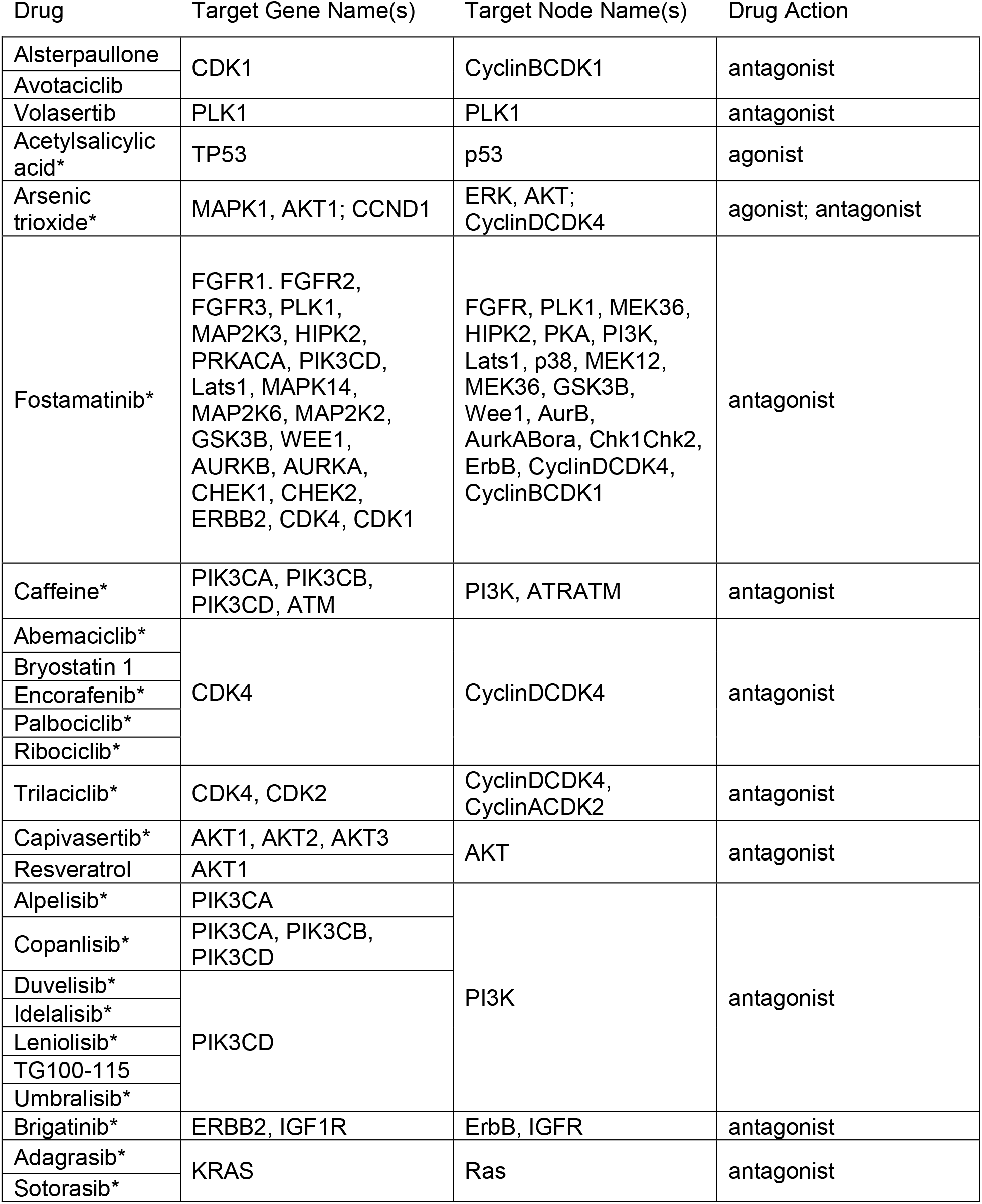
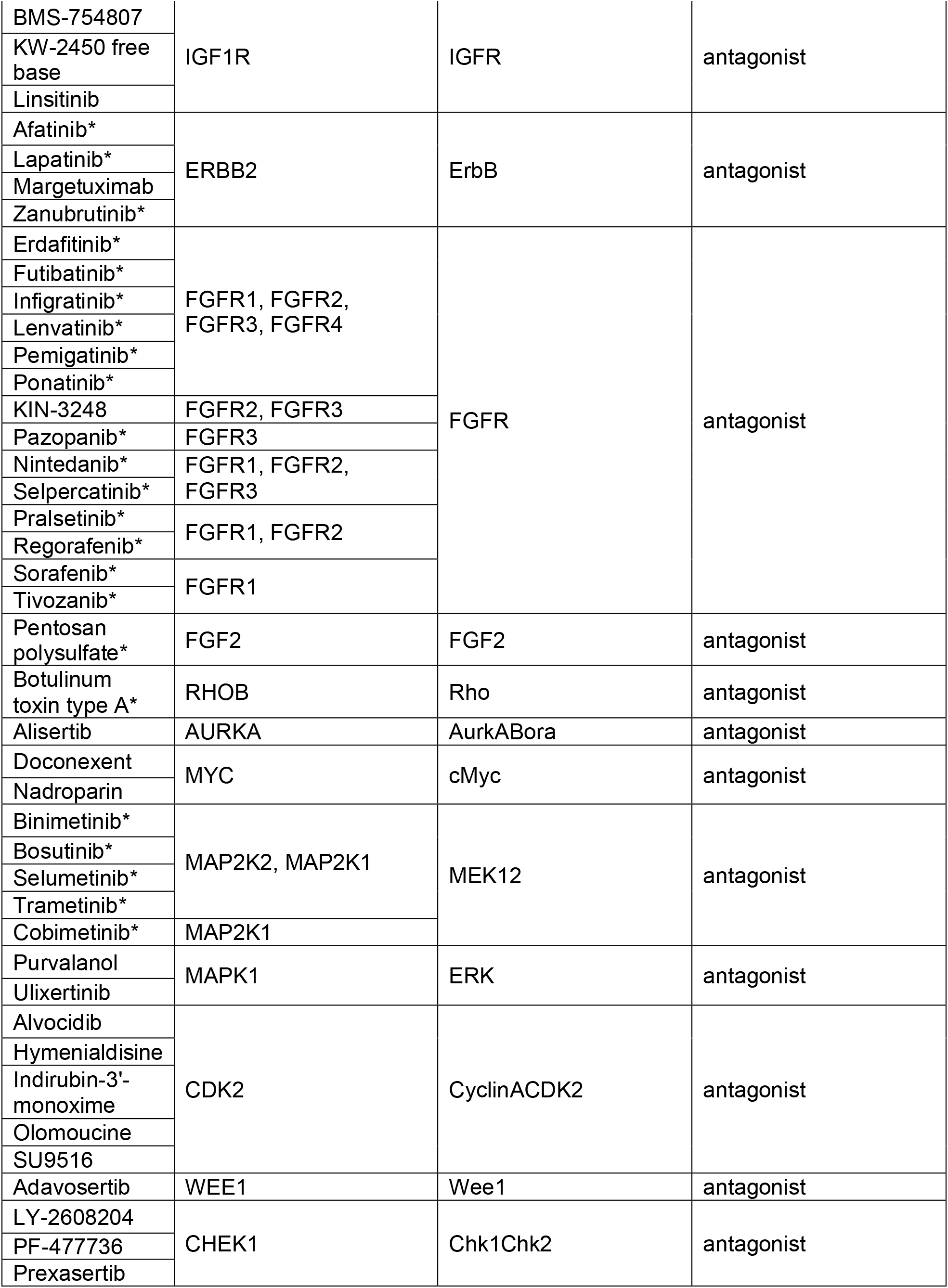
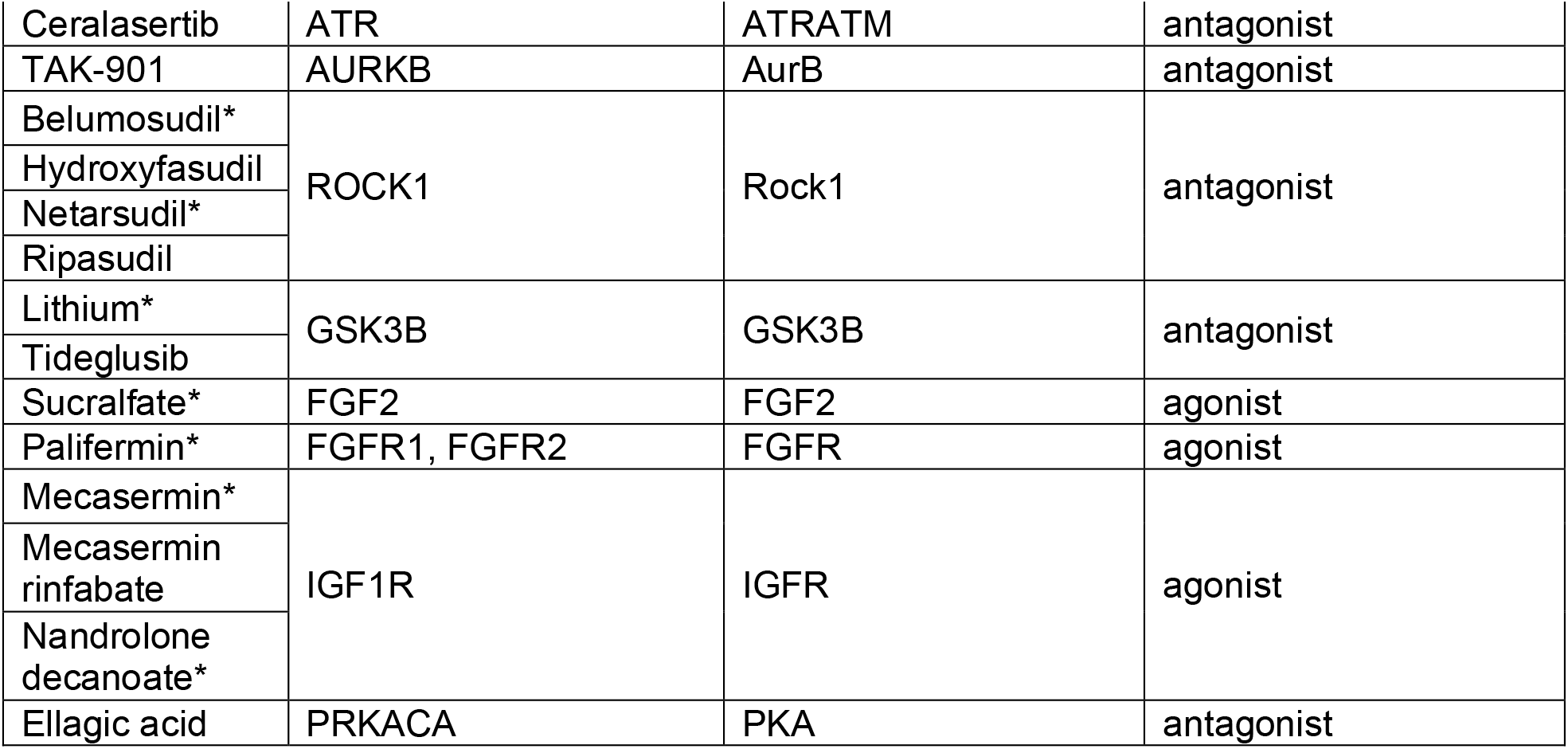
Drugs identified from the virtual drug screen as targeting the cardiomyocyte signaling network and their corresponding targets and drug action. * indicates that the drug is FDA approved.

**Supplementary Figure 1.**
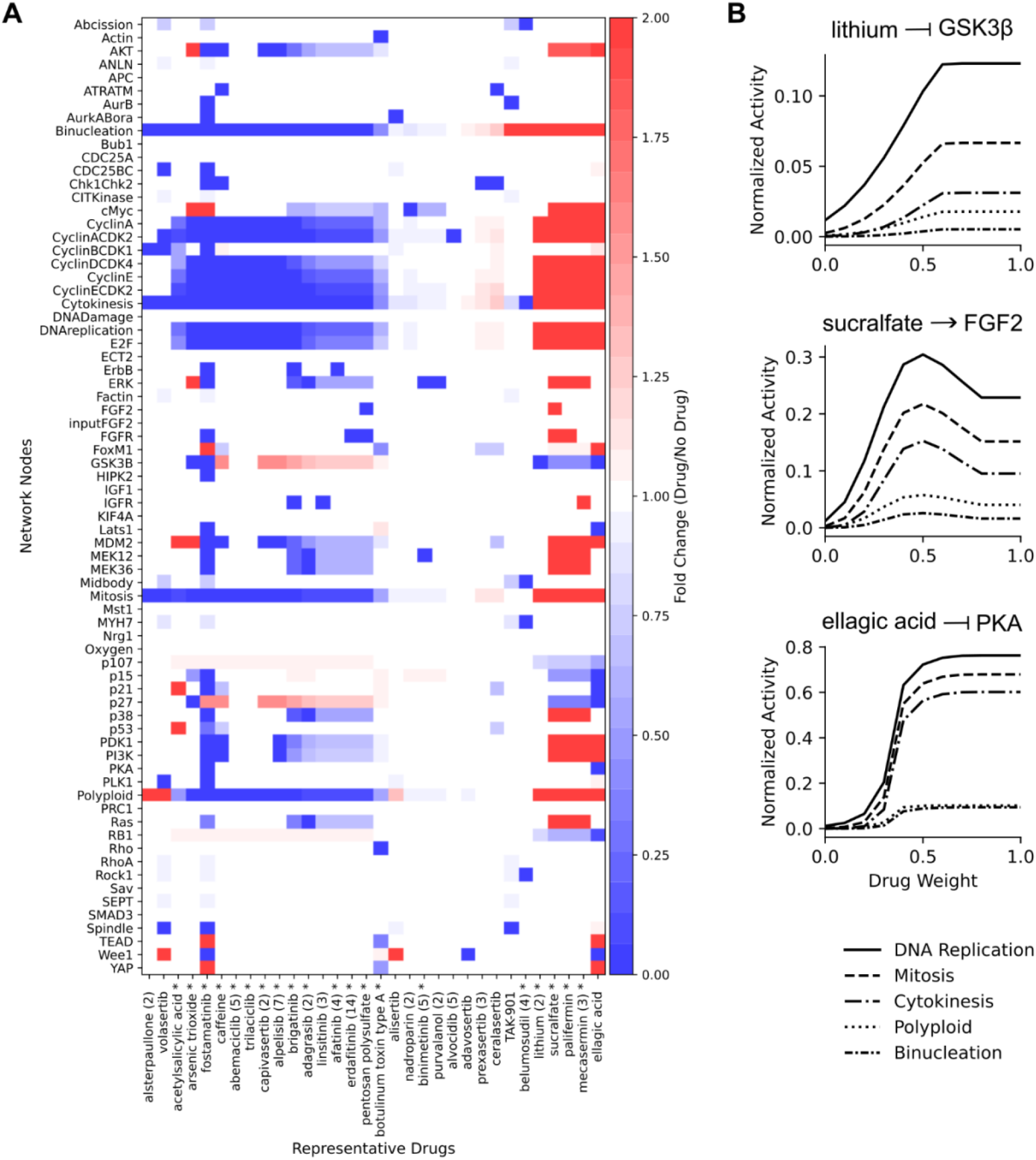
Virtual drug screen predicts that the 32 representative drugs induce distinct network states. **A)** Predicted changes in cardiomyocyte proliferation signaling network model state in response to drug simulation. (#) represents number of drugs exhibiting that unique drug target-action combination, and * indicates that the drug is FDA approved. Drugs ordered based on Figure 1B clusters. **B)** Predicted dose response for 3 pro-proliferative drugs lithium (GSK3β inhibitor), sucralfate (FGF2 agonist), ellagic acid (PKA inhibitor) exhibiting distinct targets and dose-response relationships.

### Lithium validated to increase cell cycle activity in vitro

To validate the predictions from the virtual drug screen, we tested whether the pro-proliferative drugs lithium, sucralfate, and ellagic acid increase cell cycle activity in CMs *in vitro*. Primary neonatal rat CMs were treated with selected drugs for 48 hours, and cell cycle activity was assessed by high-content microscopy of Ki67 expression and DNA content (**Figure 2A**).

**Figure 2.**
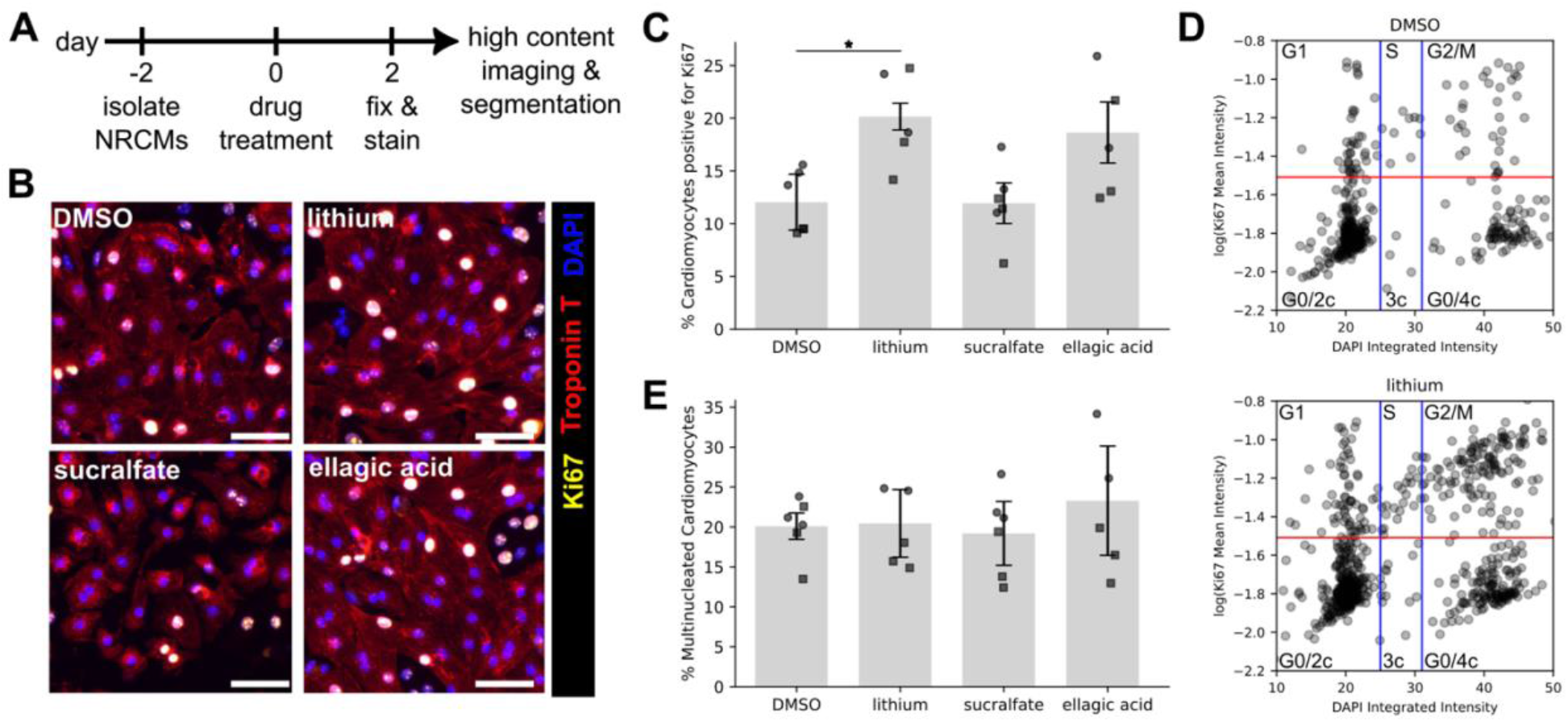
GSK3β antagonist, lithium, increases cell cycle activity in neonatal rat cardiomyocytes, as predicted by network model. **A)** Cells were treated with specified drugs for 48 hours, and cell cycle activity was measured using automated segmentation of Ki67 and DAPI. **B)** Representative images of cardiomyocytes treated with 0.1% DMSO, 10 µM lithium, 10 µM ellagic acid (PKA antagonist), or 10 µM sucralfate (FGF2 agonist). Scale bars = 100 µm. **C)** Quantification of percent of cardiomyocytes expressing Ki67. **D)** Bivariate analysis of DNA content (2c, 3c, and 4c as in blue vertical lines) and Ki67 (red horizontal line) to characterize cell cycle phase and nuclear ploidy. **E)** Percent of multinucleated cardiomyocytes. Two independent cell isolations were used and shown with circle or square data points (n = 6 wells). Two-way ANOVA with Dunnett’s post-hoc test compared to DMSO control was performed. * p<0.05.

Representative images show a qualitative increase in Ki67-positive nuclei in Troponin T-positive CMs treated with the GSK3β antagonist lithium compared to DMSO controls (**Figure 2B**). Automated image analysis of Ki67-positive CMs confirmed that lithium significantly increased cell cycle activity compared to DMSO control (**Figure 2C**). In contrast, treatment with the PKA antagonist ellagic acid or the FGF2 agonist sucralfate did not produce a statistically significant increase in Ki67 expression (**Figure 2B and C**).

To further validate cell cycle phases predicted by the network model, we performed a single-cell bivariate analysis of Ki67 expression along with DNA content, measured by DAPI integrated intensity and interpreted as 2C diploid or 4C tetraploid nuclei^25,26^. Lithium-treated CMs exhibited shifts to Ki67 positive and 4C DNA content consistent with increased G2/M cell cycle progression (**Figure 2D**). No significant differences were observed in the percentage of multinucleated CMs across treatment conditions, suggesting that these 4C CMs may be proceeding through proliferation (**Figure 2E**). Additional dose-response experiments demonstrated that lithium at 10 µM produced a saturating effect on Ki67 and Cyclin D1 expression, comparable to that observed at 1 mM lithium, the therapeutically relevant concentration approximating patient exposure^31^ (**Figure S2**). Therefore, subsequent experiments used lithium at a concentration of 10 µM. Collectively, these results indicate that lithium enhances cell cycle activity in neonatal rat CMs without altering multinucleation, consistent with promotion of later stages of cell cycle progression.

**Supplementary Figure 2.**
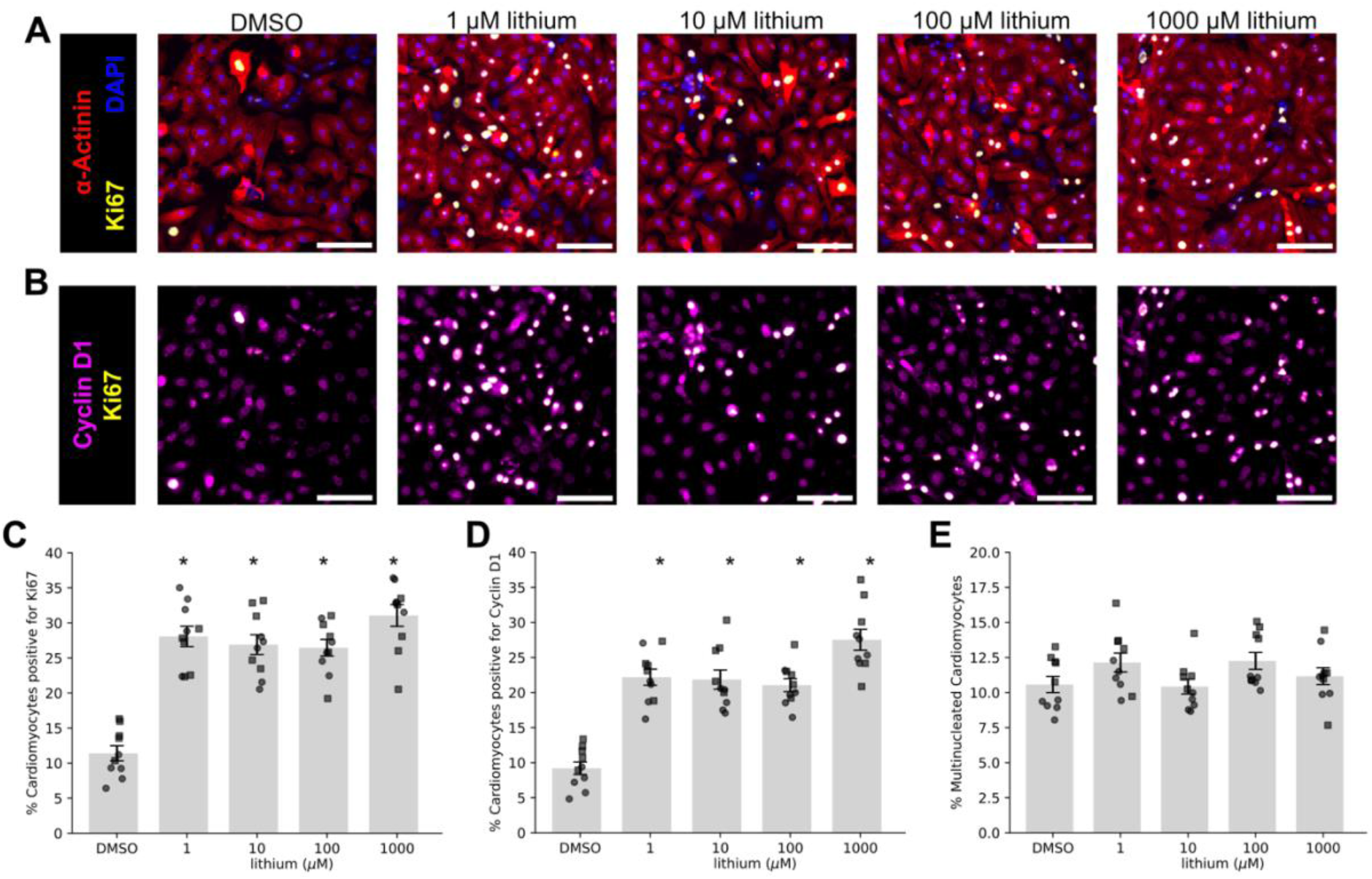
Lithium treatment increases Ki67 and Cyclin D1 expression in neonatal rat cardiomyocytes. **A-B)** Representative images of cardiomyocytes treated with DMSO or lithium (1-1000 µM). Scale bars = 100 µm. **C)** Percent of cardiomyocytes expressing Ki67 **D)** or expressing Cyclin D1. **E)** Percent of multinucleated cardiomyocytes. Two independent cell isolations were used and shown with circle or square data points (n = 10 wells). Two-way ANOVA with Tukey post-hoc test was performed. *p<0.05 compared to DMSO control.

### Cyclin D/CDK4 mediates lithium-induced cell cycle activity

To identify the molecular mechanisms by which lithium promotes CM cell cycle activity, we simulated a virtual knockdown screen in the presence of lithium. We first mapped the nodes directly perturbed by lithium treatment within the signaling network model (**Figure 3A**), then identified nodes whose knockdown modulates the effect of lithium on DNA replication (**Figure 3B**). Intersecting these two sets generated a mechanistic subnetwork^20^, highlighting the critical pathways that mediate lithium-induced DNA replication (**Figures 3C**). This analysis predicted that lithium promotes DNA replication through activation of the Cyclin D/CDK4 complex (**Figure 3D**). Network model simulations guided by the mechanistic subnetwork predicted that inhibition of GSK3β increased DNA replication in a manner dependent on Cyclin D-CDK4 activity, which could be suppressed with a CDK4 inhibitor (**Figure 3E**).

**Figure 3.**
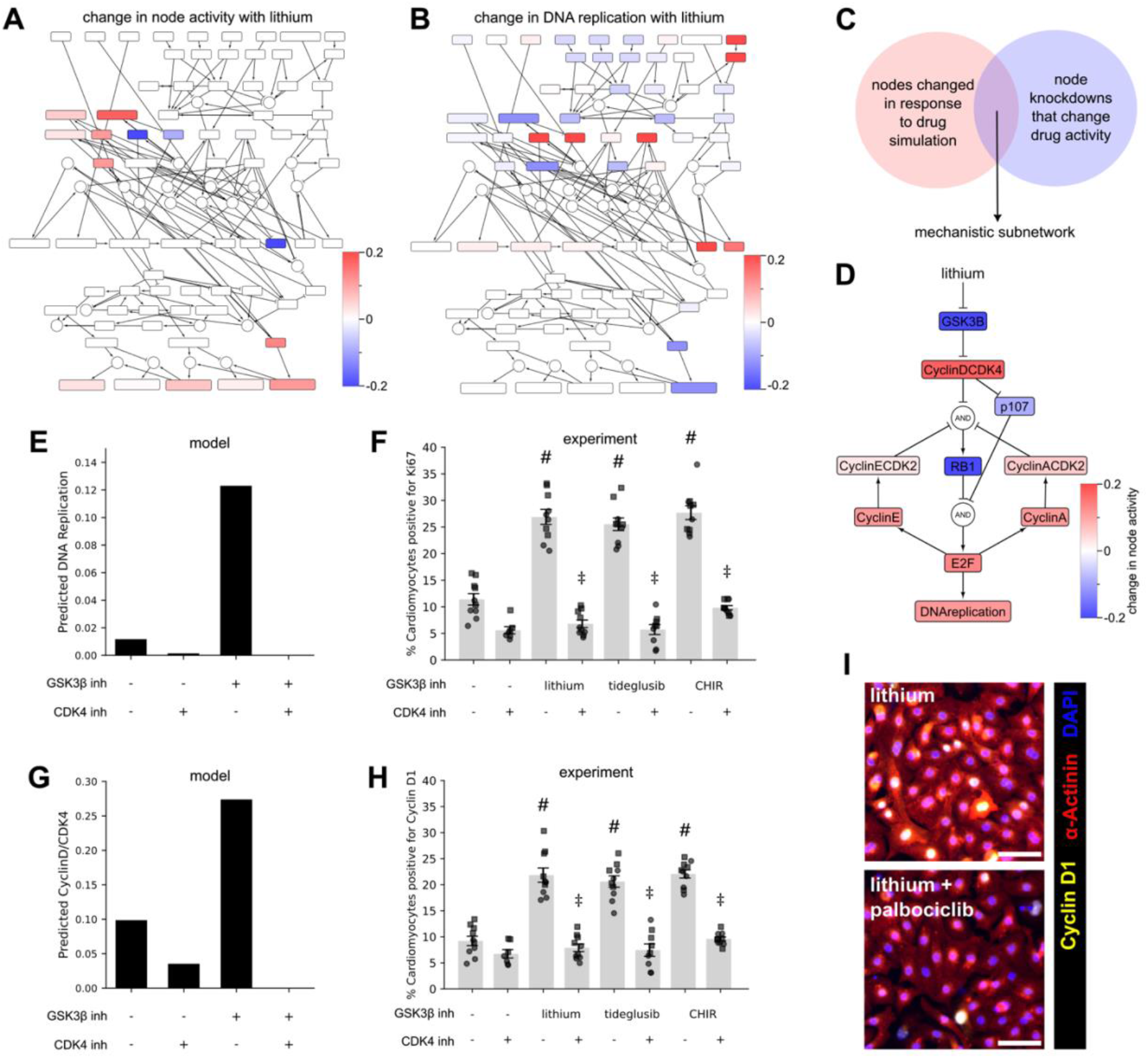
CyclinD/CDK4 mechanistically predicted and experimentally validated to mediate lithium-induced cell cycle activity in cardiomyocytes. **A)** Nodes of computational model that are altered in response to lithium. **B)** Nodes of computational model whose knockdown modulate the effect of lithium on DNA replication. **C)** A mechanistic subnetwork is identified by the intersection of nodes from panels A and B. **D)** Lithium mechanistic subnetwork identified critical nodes mediating lithium-induced DNA replication. **E)** Network model predicted effects of GSK3β inhibition and CDK4 inhibition on DNA replication. **F)** Experimental validation in neonatal rat cardiomyocytes treated with GSK3β inhibitors (10 µM lithium, 1 µM CHIR99021, or 1 µM tideglusib) and CDK4/6 inhibitor (1 µM palbociblib), measuring the percent of cardiomyocytes expressing Ki67. **G)** Network model predicted effects of GSK3β inhibition and CDK4 inhibition on CyclinD/CDK4 activity. **H)** Experimental validation in neonatal rat cardiomyocytes treated with GSK3β inhibitors and CDK4/6 inhibitor and measured percent of cardiomyocytes expressing Cyclin D1. **I)** Representative images of cardiomyocytes treated with lithium or lithium and palbociclib. Scale bars = 100 µm. Two independent cell isolations were used and shown with circle or square data points (n = 10 wells). Two-way ANOVA with Tukey post-hoc test was performed. #p<0.05 compared to DMSO control, ‡p<0.05 compared to respective GSK3β inhibitor alone.

We next experimentally tested this model-generated hypothesis, that the proliferative effect of lithium and other GSK3β inhibitors (CHIR99021 and tideglusib) depend on CDK4. Indeed, treatment with the GSK3β inhibitors lithium, CHIR99021, or tideglusib significantly increased the percentage of Ki67-positive CMs, and this response was eliminated by co-treatment with the CDK4/6 inhibitor palbociclib (FDA approved for multiple cancers) (**Figure 3F, S3**). Additional model simulations predicted that GSK3β inhibition enhances Cyclin D/CDK4 activity (**Figure 3G**). To test this prediction, we measured Cyclin D1 expression in CMs treated with GSK3β inhibitors either with or without palbociclib. Supporting the model predictions, all three GSK3β inhibitors increased the percentage of Cyclin D1-positive CMs, which was reduced by palbociclib co-treatment (**Figures 3H, I**). Together, these results demonstrate that lithium promotes CM cell cycle activity through a Cyclin D/CDK4-dependent mechanism, providing mechanistic validation of the network model predictions.

**Supplementary Figure 3.**
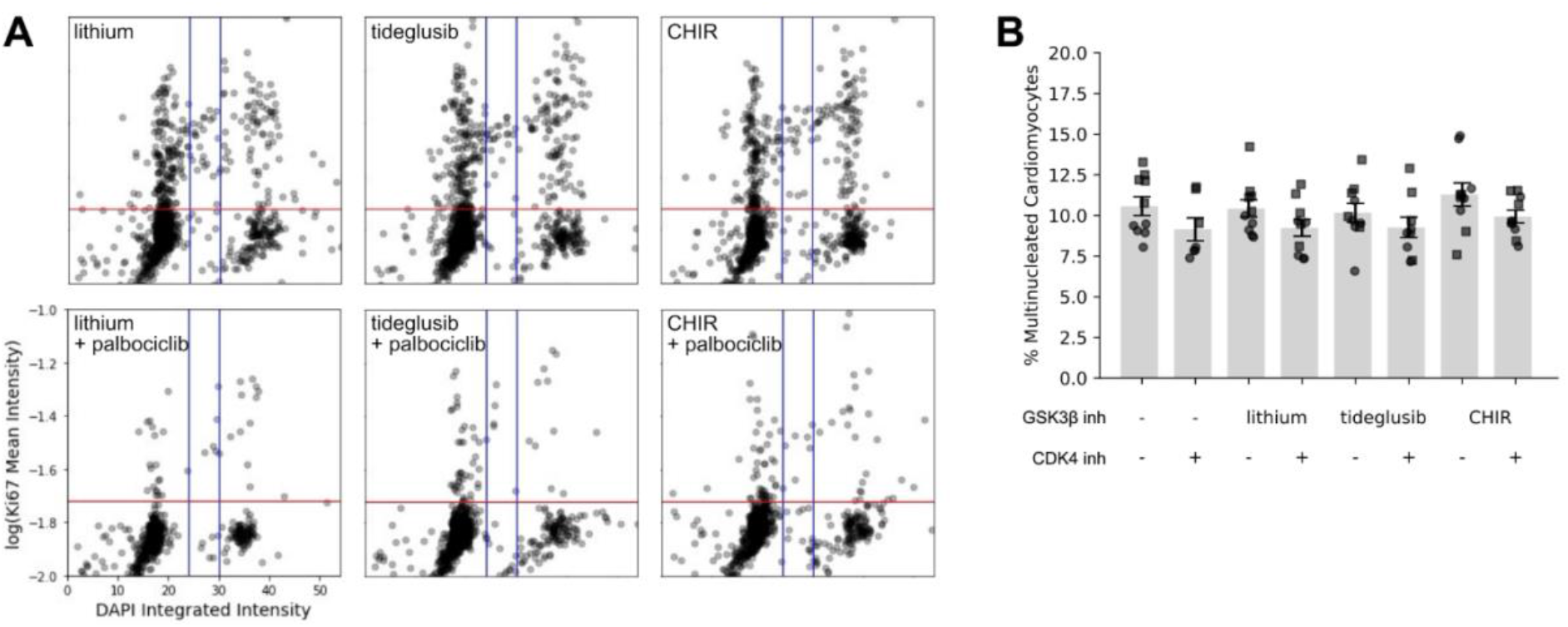
GSK3β inhibitor-mediated increase in cell cycle activity is blocked by CDK4 inhibitor palbociclib. **A)** Bivariate analysis of DNA content (2c, 3c, and 4c as in blue vertical lines) and Ki67 (red horizontal line) to characterize cell cycle phase and nuclear ploidy. Data from experiments in Figure 3. **B)** Percent of multinucleated cardiomyocytes. Two independent cell isolations were used and shown with circle or square data points (n = 10 wells). No differences observed in multinucleation using two-way ANOVA.

### Lithium validated to increase DNA replication in vivo

To determine whether lithium promotes CM proliferation *in vivo*, we evaluated the effects of maternal lithium treatment during murine cardiac development (**Figure 4A**). Pregnant C57BL/6 mice received a single intraperitoneal dose of lithium (250 mg/kg) at embryonic day (E)7.5. EdU (50 mg/kg, i.p.) was administered 4 hours before harvest at E13.5. Embryos exposed to lithium exhibited normal gross morphology as controls (**Figure 4B**). In immunofluorescence images of embryonic heart sections, lithium-exposed hearts showed more EdU incorporation within cTnT+ myocardium than saline controls (**Figure 4C**). In both groups, EdU labeling was qualitatively concentrated in the compact myocardium. To quantify these images in a consistent and scalable way, we developed an automated machine learning-based image analysis pipeline for comprehensive assessment of CM DNA replication in tissue sections (**Figure 4C**). Briefly, this pipeline uses Cellpose to segment nuclei from DAPI channel and ilastik machine-learning classification nuclei as positive or negative for cTnT and EdU. We applied this pipeline, to images from embryonic hearts, which revealed a significantly higher percentage of EdU+ CM nuclei after lithium exposure (**Figure 4D**). These data indicate that maternal lithium treatment increases CM DNA replication during cardiac development in vivo.

**Figure 4.**
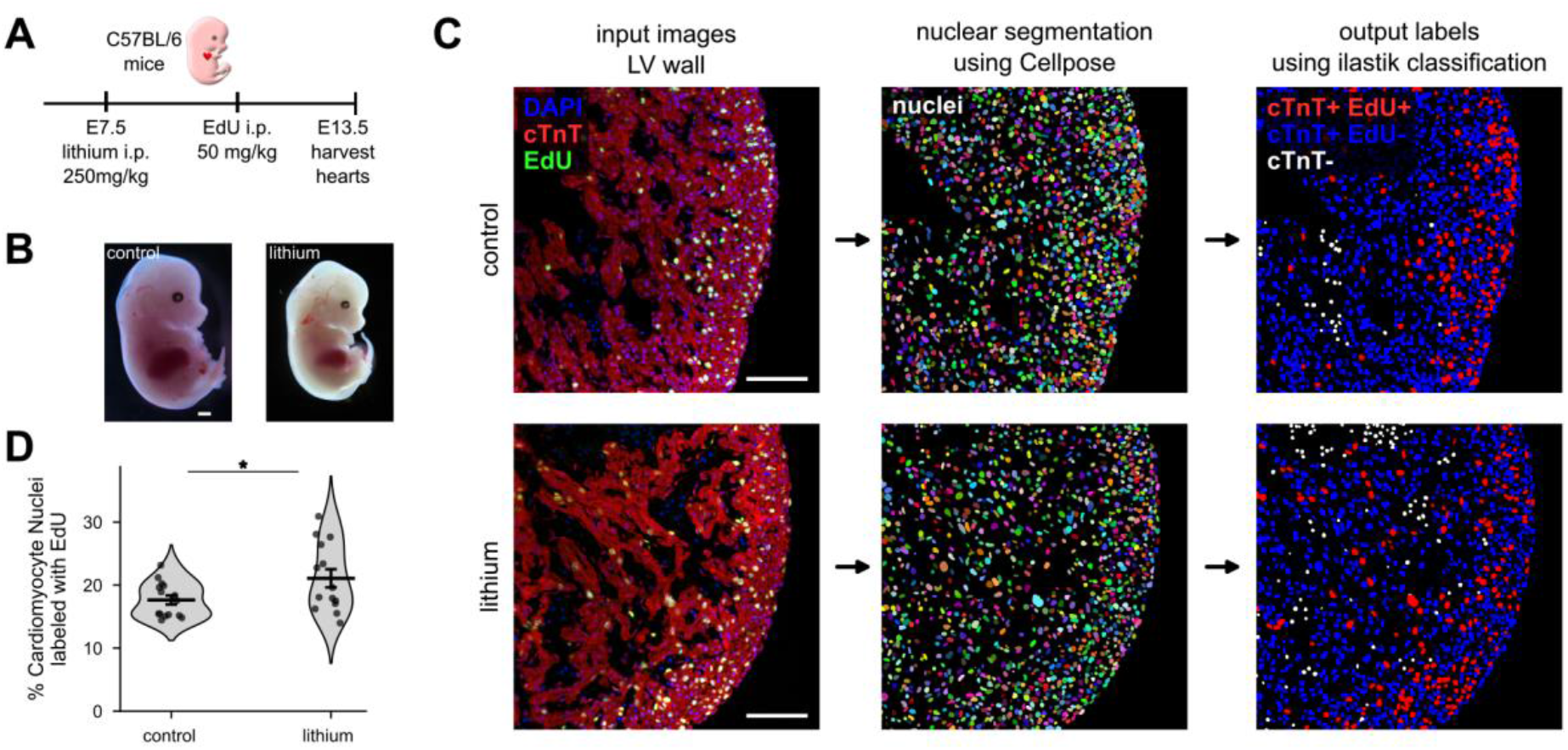
Maternal lithium treatment increased cardiomyocyte DNA replication in E13.5 embryonic hearts. **A)** Pregnant C57BL/6 mice received lithium (250 mg/kg, i.p.) at E7.5, EdU (50 mg/kg, i.p.) 4 hours before harvest, and hearts were collected at E13.5. **B)** Embryo images. Scale bars = 1 mm. **C)** Representative immunofluorescence images of embryonic heart sections stained for EdU (green), cardiac troponin T (red), and DAPI (blue). Scale bars = 100 µm. **D)** Percentage of cardiomyocyte nuclei labeled with EdU, quantified using the pipeline in C. Violin plots show the distribution with individual hearts overlaid; n = 14-15 hearts per group. *p < 0.05, two-tailed unpaired t-test.

### Patient reports support association between in utero lithium exposure and congenital heart disease

Based on the pro-proliferative effects of lithium observed in cultured rat CMs and embryonic mouse CMs *in vivo*, we next sought to determine whether *in utero* lithium exposure is associated with congenital heart disease phenotypes in humans. Lithium is widely used for bipolar disorder management during pregnancy and the postpartum period, but first-trimester exposure has been associated with an increased risk of congenital malformations^32^. More recent analyses suggest that the risk for specific outcomes, including Ebstein’s anomaly, may be smaller than previously reported^33^. However, these prior clinical studies have not been linked to mechanisms of CM proliferation, and they primarily focused on overall malformation risk and perinatal outcomes rather than specific congenital heart disease phenotypes or potential sex-specific associations.

To address these gaps, we analyzed reports from the FDA Adverse Event Monitoring System (AEMS), a pharmacovigilance database of adverse event reports on FDA-regulated products, submitted by patients, caregivers, healthcare professionals, and manufacturers. We first assessed congenital heart disease reporting across all AEMS reports. Relative to all reports in the database, reports involving lithium had higher odds of congenital heart disease (ROR = 2.01, n = 147; **Figure 5A**). This association was present in reports for males (ROR = 2.47, n = 77) but not for females (ROR = 1.04, n = 29; **Figure 5A**). We next focused on children with *in utero* exposure. Using AERSMine^29^, we identified 559,102 pediatric reports. Of these, 30,243 included cardiac-related adverse events, and 3,608 involved *in utero* exposure to a mood stabilizer (**Figure 5B**). Among all children exposed to lithium *in utero*, reporting odds were increased for three terms relative to all pediatric reports for each term: Ebstein’s anomaly, patent ductus arteriosus, and the aggregate term congenital heart disease (**Figure 5C**). When stratified by sex, male children showed increased reporting odds for patent ductus arteriosus and congenital heart disease. Female children showed increased reporting odds for patent ductus arteriosus only (**Figure 5C**). Persistent fetal circulation was reported but did not reach significance in any group.

**Figure 5.**
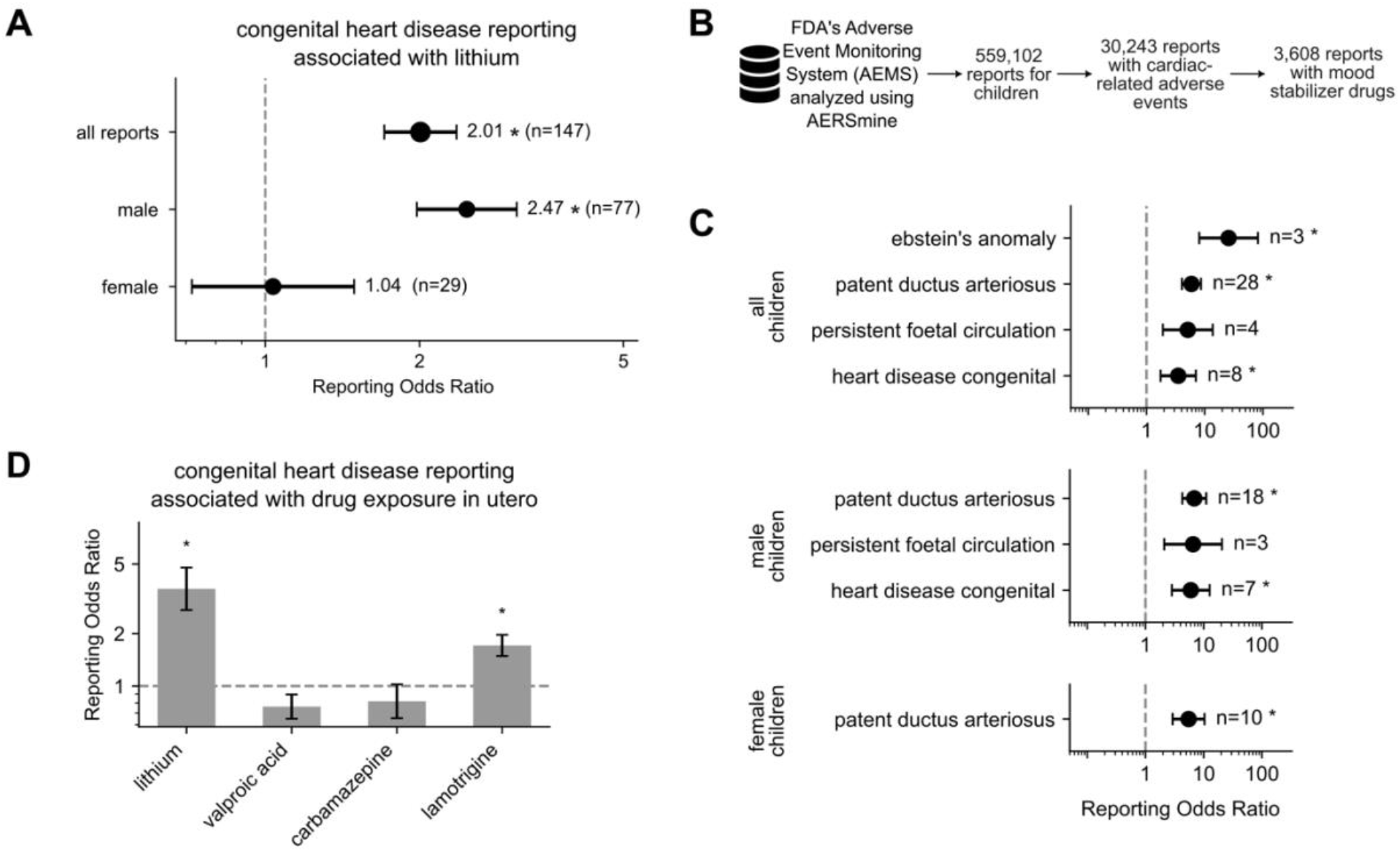
Pharmacovigilance data support an association between in utero lithium exposure and congenital heart disease. **A)** Reporting odds ratios (RORs) for congenital heart disease in all the FDA Adverse Event Reporting System (AEMS) reports involving lithium, relative to all reports in the database, shown overall and stratified by sex (sex was not reported for 41 reports). **B)** Pediatric reports in the AEMS (n = 559,102) were queried using AERSMine and filtered to reports with cardiac-related adverse events (n = 30,243) and *in utero* exposure to a mood stabilizer (n = 3,608).**C)** RORs for individual congenital cardiac adverse event terms after *in utero* lithium exposure in all, male, and female children, relative to all pediatric reports for each term. Only terms with n ≥ 3 are shown. **D)** RORs for congenital heart disease after *in utero* exposure to lithium, valproic acid, carbamazepine, or lamotrigine, relative to all pediatric reports. Error bars, 95% CI; dashed lines, ROR = 1; n, reports with both drug and event. *Fisher’s exact test, Benjamini-Hochberg FDR q < 0.05 with lower 95% CI > 1 (n ≥ 3).

To determine whether this association was specific to lithium or shared across mood stabilizers, we also examined congenital heart disease reporting after *in utero* exposure to valproic acid, carbamazepine, and lamotrigine. Because these drugs are also used to treat bipolar disorder, this comparison helps assess whether the signal reflects the drug rather than the underlying condition. Relative to all pediatric reports, lithium exposure was associated with the largest increase in congenital heart disease reporting, followed by lamotrigine. Neither valproic acid nor carbamazepine was associated with increased reporting (**Figure 5D**).

We also asked whether lithium-associated cardiac reporting extends beyond congenital phenotypes to intrinsic myocardial disease in adults. Reporting odds for intrinsic myocardial disease were increased for lithium in adults (ages 25-65) and the elderly (66+), but not in young adults (ages 15-24), where only carbamazepine was elevated (**Figure S4A**). Among lithium reports, increased reporting odds were observed for multiple specific myocardial terms in adults, including cardiomyopathy, cardiomegaly, left ventricular dysfunction, and cardiac hypertrophy (**Figure S4B**). Together, these findings indicate that *in utero* lithium exposure is associated with a distinct profile of reported congenital heart disease phenotypes, with partially sex-specific patterns and differences relative to other mood stabilizers, and that lithium exposure in adulthood is also associated with reports of myocardial disease. In the context of our experimental findings demonstrating enhanced CM cell cycle activity following lithium exposure in cells and mice, these clinical associations raise the possibility that altered CM proliferative regulation may contribute to lithium-associated congenital heart disease risk.

**Supplementary Figure 4.**
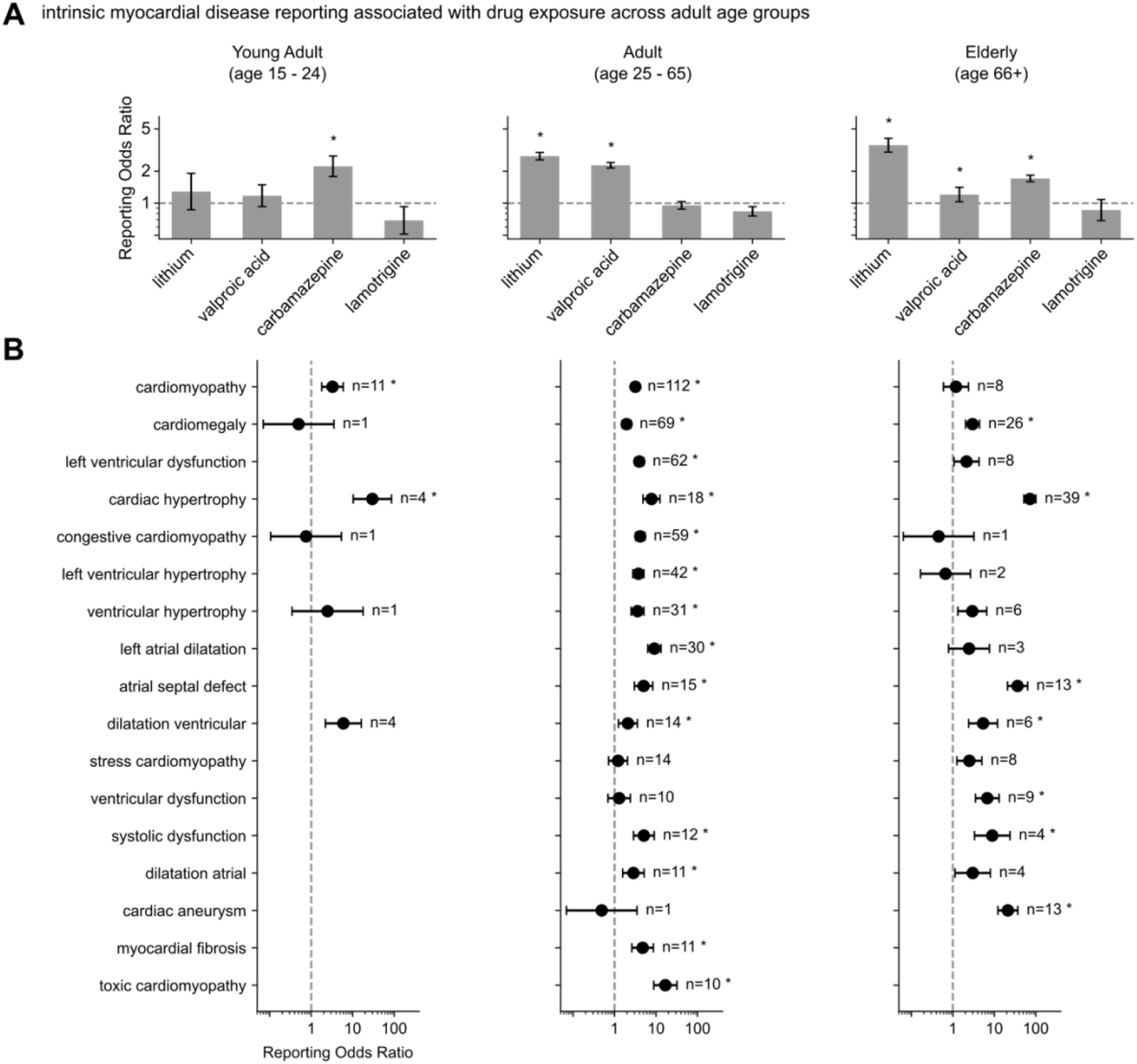
Pharmacovigilance data support an association between lithium exposure and intrinsic myocardial disease across adult age groups. **A)** Reporting odds ratios (RORs) for intrinsic myocardial disease in FDA Adverse Event Reporting System (AEMS) reports involving lithium, valproic acid, carbamazepine, or lamotrigine, relative to all adult reports in the database, stratified by age group. **B)** RORs for individual intrinsic myocardial adverse event terms after lithium exposure in young adult, adult, and elderly patients, relative to all reports for each term within the corresponding age group. Only terms with n ≥ 10 for at least one age group are shown. Error bars, 95% CI; dashed lines, ROR = 1; n, reports with both drug and event. *Fisher’s exact test, Benjamini-Hochberg FDR q < 0.05 with lower 95% CI > 1 (n ≥ 3).

## Discussion

CM proliferation is tightly regulated across developmental and adult contexts. Identifying drugs that increase or decrease CM proliferation may enable therapeutic strategies to control cardiac growth and regeneration. Here, we combined computational network modeling with experimental validation to identify FDA-approved drugs that regulate CM cell cycling. Using this computational-experimental framework, we identified lithium as a positive regulator of CM cycling and experimentally validated its pro-proliferative effects using cultured neonatal rat CMs, retrospective clinical data analysis, and *in vivo* mouse embryonic heart models. Mechanistically, lithium-dependent proliferation was mediated by Cyclin D/CDK4 signaling downstream of GSK3β inhibition. Consistent with a shared pathway mechanism, structurally distinct GSK3β inhibitors, including CHIR99021^34,35^ and tideglusib, recapitulated the pro-proliferative phenotype, supporting GSK3β inhibition as a common drug target underlying these effects.

Previous studies have demonstrated cardioprotective effects of genetic inhibition of GSK3β in a mouse model of heart failure^36^, whereas complete CM-specific loss of both GSK3 isoforms induces cell cycle dysregulation, multinucleation, and fatal dilated cardiomyopathy^37^. In parallel, pharmacologic inhibition of GSK3β using the small molecule inhibitor Neopetroside A has been shown to protect against myocardial ischemia-reperfusion injury^38^. In our studies, maternal treatment with lithium increased CM DNA replication in embryonic mouse hearts *in vivo*, suggesting that modulation of GSK3β-dependent signaling may influence cardiac growth responses during development. Furthermore, retrospective analysis of FDA Adverse Event Reporting System data identified associations between lithium exposure and specific congenital heart defects, raising the possibility that altered CM proliferative signaling may contribute to developmental cardiac phenotypes. Together, these findings suggest that lithium can modulate CM proliferation through GSK3β-dependent signaling and demonstrate the utility of the CM proliferation signaling network model for screening drug regulators of CM proliferation.

From a clinical standpoint, these findings should be interpreted as hypothesis-generating rather than as evidence of a causal risk. Lithium remains a first-line therapy for bipolar disorder, including during pregnancy and the postpartum period, where relapse after discontinuation carries substantial maternal and fetal risk^39^. Epidemiologic cohorts have reported a modest, dose-related increase in cardiac malformations after first-trimester exposure, with partially attenuated estimates in analyses that better account for confounding by indication^33^. Our pharmacovigilance analysis is consistent in direction but cannot estimate incidence, because reporting odds ratios lack exposure denominators and are subject to stimulated reporting, concomitant medications, and underlying maternal illness. Nevertheless, the specificity of the lithium signal for Ebstein’s anomaly and patent ductus arteriosus, its sex-stratified pattern, and its alignment with the pro-proliferative effects of GSK3β inhibition in embryonic hearts provide a plausible mechanistic framework that prior clinical studies lacked. Similarly, the increased reporting of cardiomyopathy, cardiac hypertrophy, and left ventricular dysfunction in adults raises the possibility that sustained GSK3β inhibition affects myocardial remodeling beyond development. This observation warrants further prospective evaluation, including whether dose, serum level, or timing of exposure modifies risk.

Several limitations of this study should be acknowledged. Our CM proliferation signaling network model^15^ has a high validation accuracy but remains incomplete, particularly with respect to regulators of multinucleation and polyploidization. Therefore, expansions of this model may be needed to more fully capture signaling mechanisms governing all CM cell cycle outcomes including bona fide proliferation. In addition, because the virtual drug screening was limited to drugs targeting nodes already within the CM proliferation network model, drugs acting through alternative or incompletely characterized pathways may have been excluded from analysis. Although the network model was extensively validated using CM experimental data^15^, the conservation of core cell cycle pathways raises the possibility that identified drugs may also influence proliferation in non-CM cell types, which will require further investigation to define potential therapeutic and adverse effects. Furthermore, while our study focused on experimentally validating pro-proliferative drug candidates, future studies should also investigate predicted anti-proliferative drugs to better understand mechanisms regulating both adaptive and maladaptive CM proliferation responses. Future work should additionally evaluate the long-term functional consequences of lithium-induced CM proliferation, including effects on CM maturation and cardiac function across developmental and adult contexts. Collectively, this study identifies lithium as a potential regulator of CM proliferation and establishes a scalable framework for systematic drug repurposing to identify regulators of CM proliferation.

## Acknowledgements

We thank Imrul Chowdhury, Claire Mueller, Mohammad Fallahi-Sichani, and Stefano Toldo for their contributions to this work.

